# Convergent evolution on *RBA1* regulates *Candidozyma auris* host adaptation

**DOI:** 10.64898/2025.12.04.692421

**Authors:** Laura A. Doorley, Alyssa A. La Bella, Sarah J. Jones, Guolei Zhao, Mohammad Asadzadeh, Quanita J. Choudhury, Suhail Ahmad, Eiman Mokaddas, Inaam Al-Obaid, Wadha Alfouzan, Khalifa Al-Banwan, Teresa R. O’Meara, Felipe H. Santiago Tirado, Ana L. Flores Mireles, Jeffrey M. Rybak

**Author notes:** Corresponding author **Jeffrey M. Rybak, PharmD, PhD** Assistant Member Department of Pharmacy and Pharmaceutical Sciences St. Jude Children’s Research Hospital 262 Danny Thomas Place Chili’s Care Center, I5304, Mail Stop #313 Memphis TN, 38105.

## Abstract

*Candidozyma auris* is an emerging healthcare-associated fungal pathogen, increasingly isolated from clinical outbreaks, with a high propensity to colonize patients and the medical environment. Here, we leveraged *C. auris* isolates from multi-facility clinical outbreaks to identify genomic patho-adaptations promoting persistence and dissemination in the healthcare environment. Genomic and phylogenetic analyses revealed loss-of-function mutations in the uncharacterized *C. auris* transcription factor gene *RBA1* to have independently emerged multiple times within these outbreaks. We demonstrate loss of *RBA1* increases *C. auris* adhesion to plastic and human keratinocytes, enhances biofilm formation, and exacerbates fungal burden in a mouse model of catheter-associated urinary tract infection. Finally, we uncover mutations in *RBA1* have emerged during multiple *C. auris* outbreaks across the globe, and that *RBA1* mutant lineages are present among large ongoing clinical outbreaks. These results reveal loss-of-function mutations in *C. auris RBA1* as a novel and clinically relevant genetic determinant of enhanced outbreak characteristics.

## INTRODUCTION

*Candidozyma auris* (formerly *Candida auris*) is an emerging pathogenic yeast first identified in 2009 and later recognized by the World Health Organization as a critical threat to global public health in 2022^1, 2^. Initially isolated from a limited number of individual patient infections, *C. auris* was soon revealed as the cause of multiple nosocomial outbreaks of invasive fungal infections across the globe^3^. Now, *C. auris* outbreaks have been reported from more than 50 countries, representing thousands of documented infections annually, including 4,514 cases in 2023 in the United States alone^4, 5, 6^. *C. auris* is distinguished from other fungal pathogens as it is not known to be ubiquitous within common natural environmental reservoirs nor is it a member of the normal human microbiota^7^. Instead, *C. auris* infections are healthcare-associated and typically occur following a lateral transmission to vulnerable patients within hospital settings^8^. Multiple studies have demonstrated that once introduced into the medical environment, *C. auris* rapidly spreads among both patients and medical surfaces, acting as an invasive species in these healthcare settings. As a result, clinical *C. auris* infections routinely occur in series or as clustered clinical outbreaks, sometimes spanning multiple facilities or institutions within an affected healthcare network^9, 10^.

As with other healthcare-associated and multi-drug-resistant pathogens, the characterization and tracking of emergent microbial lineages with traits impacting clinical outcomes is paramount to infection prevention and treatment. Prior study of *C. auris* evolution during infection and exposure to clinically relevant stressors has largely focused on the emergence of antifungal resistance^11, 12, 13, 14^. We and others have shown, through both *in vitro* evolution experiments and by studying sets of isogenic clinical isolates, that *C. auris* is capable of rapidly adapting to antifungal-induced stress and that multiple independent drug-resistant lineages can emerge after only a single antifungal exposure^11, 12, 13, 14^. However, investigations of patho-adaptation in the clinical setting or during outbreaks of infections remains unexplored.

In this work, we leveraged a collection of *C. auris* clinical isolates from multi-institutional outbreaks of infection to identify novel genetic markers and link them to traits driving host adaptation and outbreak potential in clinical settings. We identified multiple independent lineages of *C. auris* harboring distinct mutations in a previously uncharacterized fungal transcription factor, here named, Repressor of Biofilm and Adhesion 1 (*RBA1*). Genetic and transcriptomic interrogation revealed loss of *RBA1* to result in enhanced biofilm formation, increased adhesion to multiple surfaces including human keratinocytes, and significantly elevated expression of *C. auris* adhesins and virulence factors. Moreover, loss of *RBA1* was found to significantly increase fungal burden in a mouse model of catheter-associated urinary tract infection (CAUTI), an infection characteristically involving biofilms in urine where *C. auris* is often isolated^15^. Finally, analysis of publicly available whole genome sequencing data revealed numerous distinct *RBA1* mutations present in hundreds of clinical isolates, demonstrating consistent and convergent loss-of-function mutations from isolates from well-documented and ongoing clinical outbreaks of *C. auris* infections across the globe. Collectively these findings demonstrate *RBA1* encodes a repressor of *C. auris* biofilm and adhesion and implicate mutations in *RBA1* as clinically relevant adaptations promoting *C. auris* survival during healthcare-associated outbreaks of infection.

## RESULTS

### Clinical emergence of multiple *C. auris* outbreak lineages carrying *RBA1* mutations

Whole genome sequencing (WGS) analysis revealed all *C. auris* isolates belonged to Clade I, one of three major genetic clades previously linked to healthcare-associated outbreaks^16^. This included 34 and 36 isolates from the previously described subclades Ib and Ic, respectively^17^. A total of 155,135 SNPs were identified among the 70 included isolates as compared to the Clade I reference genome (B8441v3), with 46,231 predicted to impact coding regions. As anticipated from characteristically fluconazole-resistant Clade I isolates, all isolates were found to harbor one or more mutations in the genes *ERG11*, *TAC1B*, and/or *MRR1*, which we and others have previously shown to directly contribute to fluconazole resistance^14, 18, 19, 20^. Additionally, 16 isolates were found to possess mutations known to contribute to echinocandin resistance (altering *FKS1*) and 4 isolates possessed mutations in genes associated with amphotericin B (AmB) resistance (*ERG3*, 1 isolate; *ERG6*, 3 isolates)^12, 21, 22^.

Phylogenetic analysis revealed the population structure of this collection to consist of 69 isolates grouping into three major clusters, with 35, 24, and 10 isolates grouping into clusters A, B, and C, respectively (**Fig. 1**). A single isolate (SKU043) did not group into a larger cluster and was more distantly related. Within clusters, isolates were found to be largely clonal, with an average of 5.3 to 12.6 coding region SNPs separating each isolate (**Fig. S1**), while SKU043 differed from all other isolates by 43 to 113 coding region SNPs. Clusters A and C each contained isolates originating from multiple healthcare facilities within Kuwait, while cluster B originated entirely from a single tertiary care hospital. Notably, multiple instances were identified where isolates from different patients and even different hospitals were found to be more genetically similar than serial isolates originating from the same patient case, as might be expected in the setting of a clinical outbreak.

**Fig 1.**
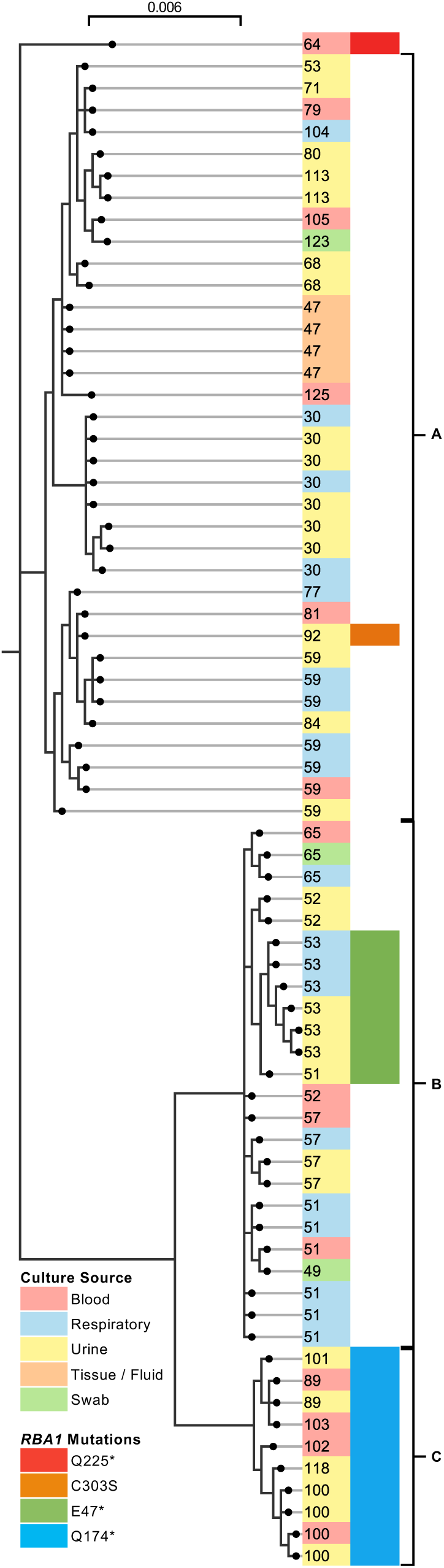
Clinical emergence and spread of *C. auris* isolates harboring mutations in *RBA1* during an outbreak of infections. Phylogenetic analysis of 70 isolates from documented outbreaks of *C. auris* infections in Kuwait. Nodes indicate branch length termination. Patient cases, culture sources, and *RBA1* mutations are indicated on the right.

To investigate potential patho-adaptations occurring during outbreaks of *C. auris* infections, we further evaluated the WGS data to identify genes where mutations were acquired on multiple occasions and/or mutations emerged and persisted among the outbreak population. Intriguingly, this revealed 19 isolates harboring mutations in a previously uncharacterized *C. auris* gene B9J08_03209 (identified as B9J08_002136 in B8441v2 and hereafter named *RBA1*), which is predicted to encode a fungal-specific zinc-cluster transcription factor (ZCF). Although initially identified as an ortholog of *WOR2* in *C. albicans,* the proteins are not syntenic, with only 22% sequence identity, primarily located in the DNA-binding domain. The 19 isolates with mutations in *RBA1* included representatives from all three major phylogenetic clusters, including all 10 isolates in cluster C, as well as the non-clustered isolate SKU043. Four different mutations in *RBA1* were identified, all considered likely loss-of-function mutations, with three encoding an early stop codon (upstream of the predicted DNA-binding domain) and one altering a highly conserved cysteine residue in the Zn(II)_2_Cys_6_ DNA-binding motif. The early stop codon mutation, *RBA1*^E^^47^*, was found to have emerged and persisted among two intensive care unit (ICU) patients, these cases and the associated isolates were further evaluated to identify the potential clinical impact of *RBA1* mutations.

### Loss of *RBA1* enhances *C. auris* biofilm formation

A subset of isolates from patient cases 51 and 53 have been previously characterized with consideration to antifungal susceptibility while investigating the emergence of echinocandin and amphotericin B resistance-conferring mutations, respectively^12, 23, 24^. In brief, both were adult patients with complex medical histories including hematologic malignancies admitted to the ICU of a tertiary care center and received multiple courses of the antifungals liposomal amphotericin B (AmB) and caspofungin (CSF). Over the course of approximately eight months, a total of 14 *C. auris* isolates were cultured from the respiratory tract, urinary tract, and blood of patients 51 and 53 (**Fig. 2a**). Notably, while the earliest isolate in these cases, SKU013, harbored a wildtype *RBA1* allele (*RBA1*^WT^), an early stop codon encoding mutation in *RBA1* (*RBA1*^E47*^) emerged in an isolate obtained from culturing the urinary catheter (SKU018). This mutation was subsequently found in all but one *C. auris* isolate cultured from patient 53. Further details of these cases can be found in **Supplemental Data**

**Fig 2.**
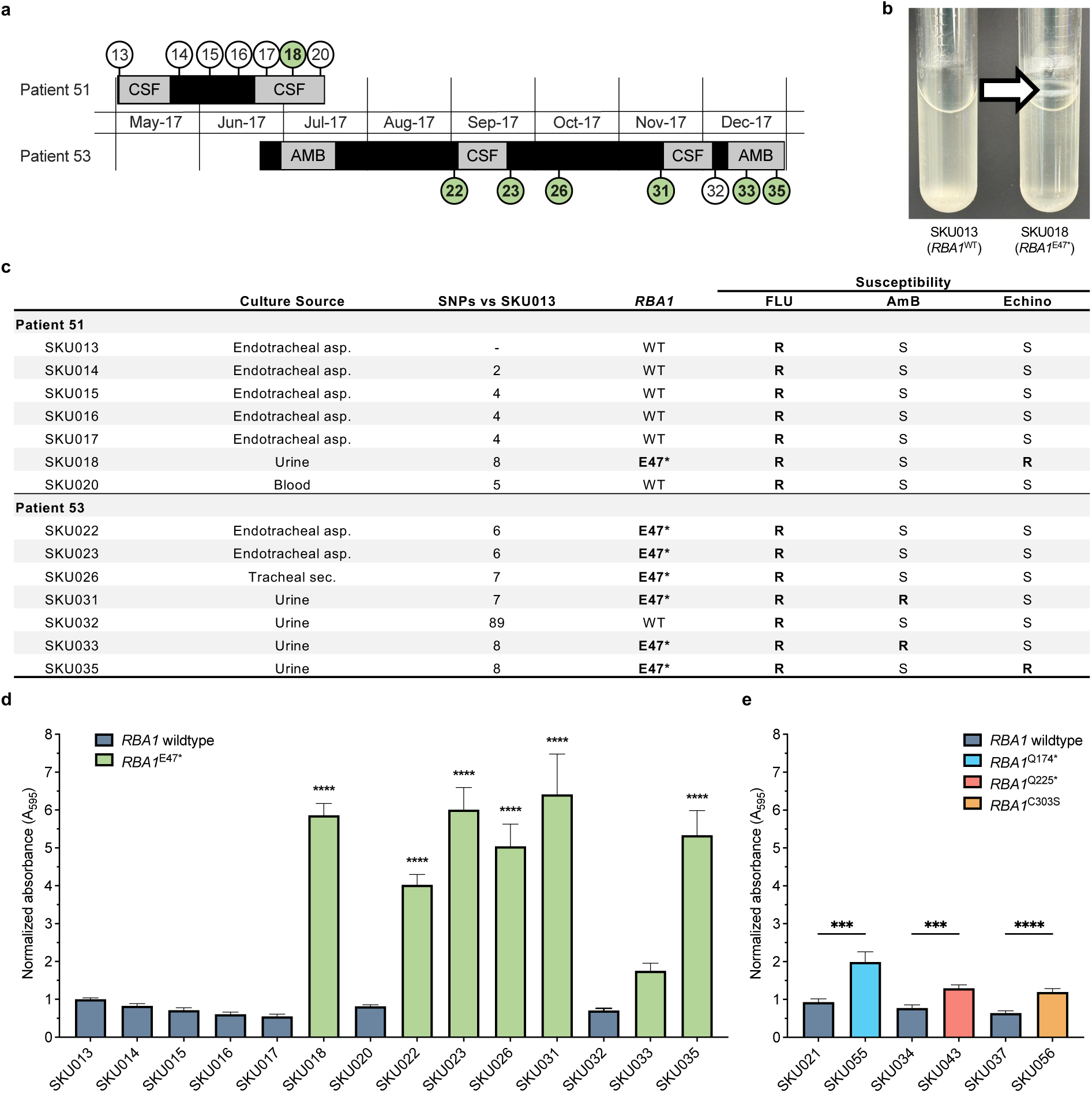
*RBA1* mutations emerge during patient infection and are associated with increased biofilm production. **a**, Timeline of patient cases 51 and 53 with *C. auris* isolations and antifungal treatment courses indicated, AMB: liposomal amphotericin B, CSF: caspofungin. Isolate numbers (SKU) with *RBA1^E^*^47^*\**mutations indicated by green background. **b,** Biofilm formation on sidewall of polystyrene (PS) tubes following overnight growth in RPMI culture media. Arrow indicates visible biofilm ring. **c,** Isolate details including name, culture source, number of coding region single nucleotide polymorphisms (SNPs) relative to the earliest isolate SKU013, *RBA1* genotype, and interpretation of susceptibility to fluconazole (FLU), amphotericin B (AmB), and echinocandins (Echino) based on current CDC tentative breakpoints. **d, e,** Biofilm production of isolates formed over 48 hours on fibrinogen-treated PS in urine media supplemented with 10% serum. Biofilm assessment via crystal violet staining with absorbance read at 595 nm. Reads were normalized to the initial isolate SKU013 with means (bars) and standard error of the means plotted in GraphPad Prism. (**d**) gray-blue bars indicate wildtype *RBA1* containing isolates and green indicates *RBA1*^E47*^ mutation. Clinical isolates harboring (**e**) *RBA1*^Q174*^, *RBA1*^Q225*^, and *RBA1*^C303S^ mutations are indicated with light blue, red and orange bars, respectively, and plotted alongside a corresponding closest genetic relative harboring a wildtype *RBA1* (gray-blue bars). Kruskal-Wallis statistical one-way ANOVA testing with Dunn’s correction for multiple comparisons showed significant differences between SKU013 and *RBA1*^E47*^ containing isolates (**d**; \*\*\*\**P* = <0.0001) and *RBA1*^Q174*^, *RBA1*^Q225*^, and *RBA1*^C303S^ isolates and corresponding closest genetic relative (**e**; \*\*\**P*= <0.001).

Of the 14 isolates cultured from patient 51 and patient 53, 13 were found to be closely related (separated by 2 to 8 SNPs within coding regions, **Fig. S1**) and grouped into cluster B. While *RBA1* mutant isolates were observed to exhibit increased biofilm formation under standard laboratory conditions, there was no association between antifungal susceptibility and *RBA1* genotype, (**Fig. 2b, c**). As the earliest isolate where the *RBA1*^E47*^ mutation was present (SKU018) was cultured from the urinary tract of patient 51, and multiple subsequent isolates with this mutation were isolated from the urinary tract of patient 53, we used our fibrinogen-urine biofilm model ^25^ that utilizes serum-supplemented urine and fibrinogen-coated surfaces to evaluate biofilm formation in conditions mimicking the bladder during CAUTI. In this model, isolates carrying the *RBA1*^E47*^ mutation exhibited enhanced biofilm formation as compared to the earliest isolate SKU013 (approximately 2 to 6-fold higher), while biofilm formation was relatively unchanged among closely related *RBA1*^WT^ isolates (**Fig. 2d**). To evaluate if other clinically emergent predicted loss-of-function mutations in *RBA1* had a similar effect on biofilm formation, we also compared the biofilm formation of isolates harboring the *RBA1*^Q174*^ (SKU055), *RBA1*^Q225*^ (SKU043), and *RBA1*^C303S^ (SKU056) mutations to their closest available genetic relatives (SKU021, SKU034, and SKU037, respectively) using the same model. In each case, biofilm formation was observed to be significantly higher in the isolates harboring mutations in *RBA1* (**Fig. 2e**), further bolstering the connection between loss of *RBA1* and biofilm formation.

From these observations, we hypothesized that loss-of-function mutations in *RBA1* are a genetic determinant of enhanced *C. auris* biofilm formation. To test this hypothesis, we next generated the *RBA1*-knockout (Δ*rba1*) and *RBA1*-revertant (Δ*rba1*::*RBA1^+^*) control strains in the SKU013 clinical isolate background and evaluated biofilm formation. In the fibrinogen-urine model, Δ*rba1* strains were found to exhibit significantly increased biofilm formation (3.9 to 4.8-fold greater than SKU013; p<0.0001). This enhancement of biofilm formation resembled the isogenic clinical isolate SKU018 which carries the *RBA1*^E47*^ mutation (3.3-fold greater than SKU013; p<0.0001) (**Fig. 3a**). Importantly, restoration of the *RBA1* gene in Δ*rba1*::*RBA1^+^*control strains returned biofilm formation to levels comparable to SKU013.

**Fig 3.**
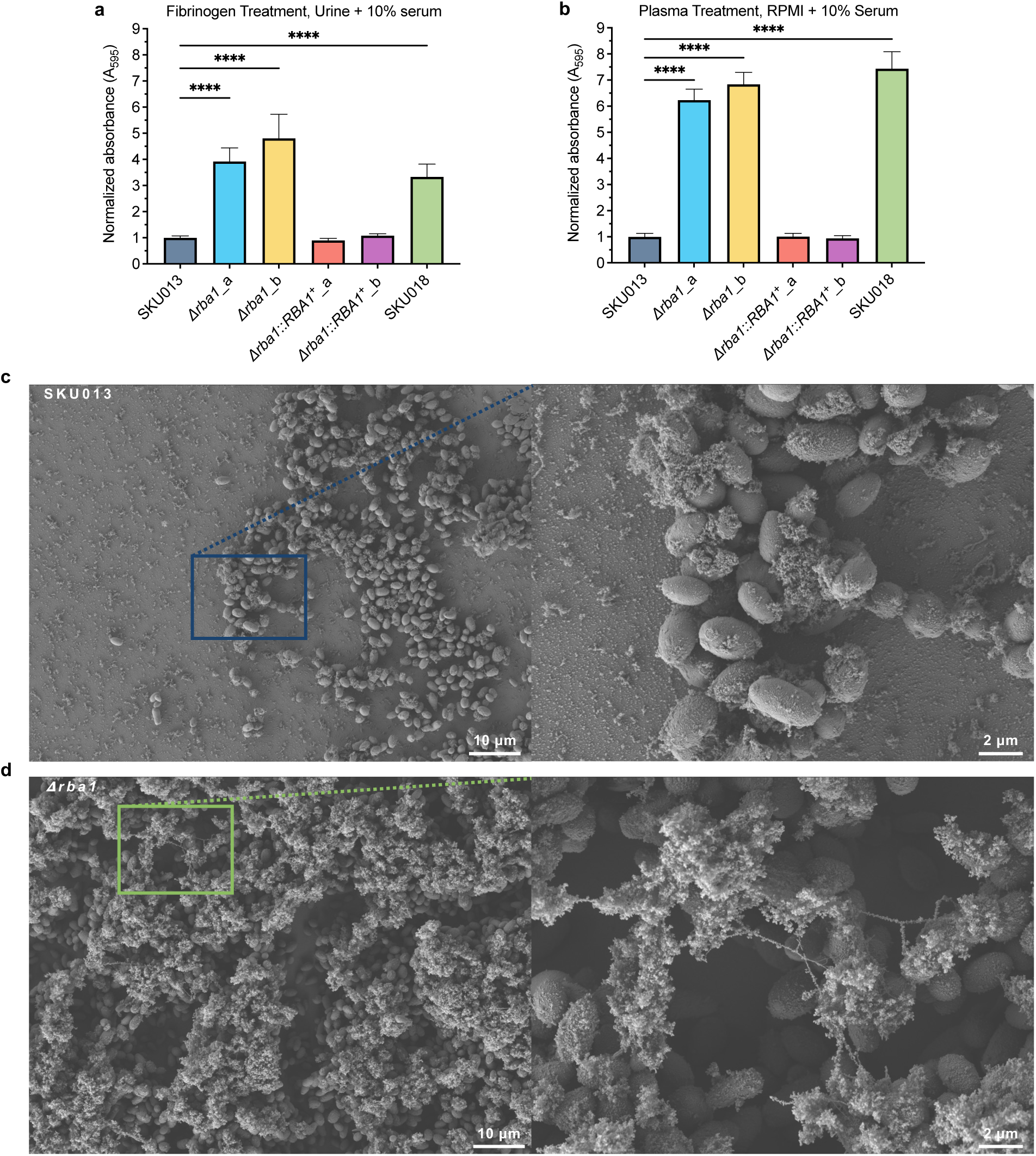
Loss of *RBA1* enhances biofilm production. **a,b,** Biofilm production of *C. auris RBA1* strains formed on (**a**) fibrinogen coated microtiter plates in human urine media supplemented with 10% human serum after 48hrs and (**b**) on human plasma coated microtiter plates in RPMI tissue culture media supplemented with 10% human serum after 24hrs. Biofilms stained with crystal violet and absorbance measured at 595 nm. Absorbance reads were normalized to SKU013 (*RBA1*^WT^) and graphed (GraphPad Prism) showing mean (bars) and standard error of the mean. Kruskal-Wallis statistical testing with Dunn’s correction for multiple comparisons showed significant differences between Δ*rba1_a,* Δ*rba1_b*, and SKU018 (*RBA1*^E47*^) when compared to SKU013 (*RBA1*^WT^) (\*\*\*\**P*= <0.0001). **c, d,** Representative SEM images for (**c**) SKU013 (*RBA1*^WT^) and (**d**) Δ*rba1* biofilms formed over 24 hours in RPMI media supplemented with 10% human serum on silicone disks treated with human plasma. Dried samples coated with 5nm iridium and imaged in the Zeiss Gemini 460 (CITC-SJCRH) using Secondary Electrons detector at 2kV, 77pA, and 5mm working distance.

We further investigated the impact of *RBA1* using a plasma-RPMI biofilm model that utilized serum-supplemented RPMI culture media and human plasma-coated surfaces to evaluate biofilm production that could apply more generally to clinical settings. Once again, Δ*rba1* strains were found to produce more robust biofilms (6.2 to 6.8-fold greater than SKU013; p<0.0001), while Δ*rba1*::*RBA1^+^*control strains closely matched the biofilm formation of SKU013 (**Fig. 3b**). Examination of the plasma-RPMI biofilms using scanning electron microscopy, showed scattered patches of biofilm predominantly consisting of one to two layers of cells for both SKU013 and Δ*rba1*::*RBA1^+^* (**Fig. 3c**). By comparison, Δ*rba1* strains formed more confluent biofilms of variable but generally greater thickness with greater production of extracellular matrix. (**Fig. 3d**).

### Loss of *RBA1* increases expression of *C. auris* adhesins and virulence factors

As *RBA1* is predicted to encode a transcription factor and our earlier experiments had shown loss of *RBA1* significantly enhances biofilm formation, we next evaluated the impact of *RBA1* on the *C. auris* transcriptome during both planktonic and biofilm growth conditions using our Δ*rba1* strain and the parental *RBA1*^WT^ isolate SKU013. When cells were grown in RPMI tissue culture media under planktonic conditions, transcriptional profiling revealed loss of *RBA1* to have a modest impact, and only 55 of 5,594 *C. auris* genes (0.98%) were differentially expressed (≥|1.5| fold-change and false discovery rate [FDR] ≤0.05) (**Fig. 4a**). Of these, 42 genes were upregulated in the absence of *RBA1*, and these genes were enriched for those involved in biofilm formation (p= 1.60e-07) and cell aggregation (p= 2.18e-07), and no enrichment for biological processes was found among the 13 down-regulated genes. By comparison, when cells were harvested from actively growing biofilms, 701 of 5,594 genes (12.53%) were observed to be differentially expressed, including 250 and 451 genes which were up- and down-regulated in the Δ*rba1* strain, respectively (**Fig. 4b**). Up-regulated genes were enriched for those involved in transmembrane transport (p= 0.01351), regulation of RNA biosynthetic process (p=0.02583), cell aggregation (p=0.05187), and regulation of primary metabolic process (p=0.07722), while down regulated genes were enriched for 18 different biological processes including protein biosynthetic process (p= 2.35e-28), macromolecule biosynthetic process (p= 4.05e-10), premeiotic DNA replication (p= 0.00273), and steroid biosynthetic process (p= 0.02579). Thirty-two *C. auris* genes were up-regulated in the absence of *RBA1* under both growth conditions, and these genes were enriched for those involved in biofilm formation and cell aggregation (**Fig. 4c**). Notably, this included four *C. auris* adhesins (*ALS4112/ALS5, IFF4*, *IFF4109*, and *SCF1*), three of which we have previously shown to impact virulence in a mouse model of invasive candidiasis^26, 27^.

**Fig 4.**
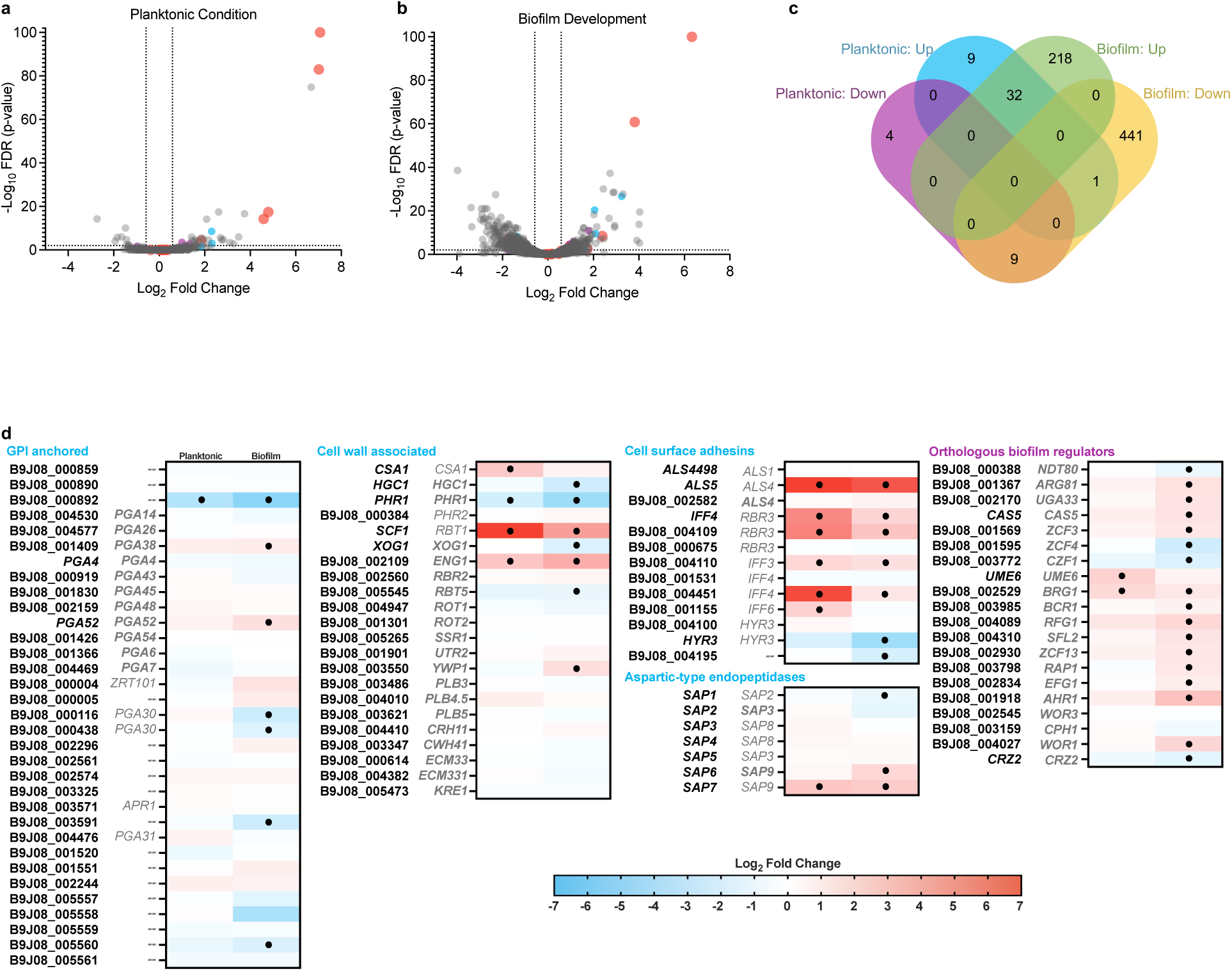
Loss of *RBA1* increases expression of *C. auris* adhesins and virulence factors under both planktonic and biofilm conditions. **a,b** Volcano plot of differentially expressed genes between clinical isolate SKU013 and *Δrba1* strain during exponential growth phase when cultured under (a) planktonic growth conditions (10mL RPMI media in 50mL conical tube incubated at 35°C with 220RPM) and (**b**) biofilm-forming conditions (10mL RPMI media in plasma-treated T75 flasks for static incubation at 37°C). Blue, purple, and red highlighted dots represent cell wall associated, ZCF, and cell aggregation or biofilm formation encoding genes, respectively. Dotted lines represent lower limit equivalents for fold changes of ≥ |1.5| on the x-axis, and a false discovery rate (FDR) p-value ≤0.05 on the y-axis. **c**, Venn diagram comparing differentially expressed genes (≥ |1.5| on the x-axis, and a false discovery rate p-value ≤0.05) exhibiting up-regulation or down-regulation in the *Δrba1* strain under planktonic or biofilm growth conditions. **d**, Differential gene expression encoding predicted GPI-anchored proteins, cell surface adhesions, fungal cell wall associated proteins, aspartic-type endopeptidases, and genes orthologous to transcriptional regulators of biofilm formation in *Candida albicans* with planktonic expression on the left and biofilm expression on the right for each column. *C. albicans* genes are listed in gray with true orthologs bolded. Significant differential expression (fold change ≥ |1.5|; FDR p-value ≤ 0.05) is indicated with a black dot.

As many genes in the *C. auris* genome remain uncharacterized and/or currently have incomplete annotations, we further evaluated differential expression among *C. auris* genes with homology to those associated with adhesion, biofilm formation, and cell wall architecture in the model pathogenic yeast *Candida albicans.* This revealed additional *C. auris*-specific adhesins, cell wall-associated proteins, GPI-anchored proteins, and aspartic-type endopeptidases exhibiting differential expression under biofilm conditions in the absence of *RBA1* (**Fig. 4d**). Notably, four of the differentially expressed genes associated with the fungal cell wall, B9J08_002109, B9J08_003350, *PHR1*, and *XOG1*, are the predicted orthologs of *C. albicans* cell wall proteins known to be involved in cell wall remodeling, cell wall masking from immune detection, and response to host immune cells^28, 29, 30^.

As *RBA1* appears to repress biofilm formation in *C. auris,* we also examined the expression of *C. auris* orthologs of genes known to regulate biofilm formation in the model pathogenic yeast, *C. albicans* (**Fig. 4d**)*. UME6* and B9J08_002529 (ortholog of *C. albicans BRG1*), exhibited significantly increased expression in the Δ*rba1* strain under planktonic conditions. In *C. albicans,* these genes are known to be important positive regulators of hyphal growth and biofilm formation. Further, *UME6* has previously been shown to impact *C. auris* biofilm formation when overexpressed and artificially hyper-activated through HA-tagging of the putative transcription factor activation domain^31^. Under biofilm conditions, 14 genes orthologous to those known to regulate biofilm formation in *C. albicans* were up-regulated, notably again including the *C. auris* orthologs of *BRG1.* These findings demonstrate that *RBA1* significantly impacts regulation of *C. auris* gene expression during biofilm formation and suggest that *RBA1* may play a central regulatory role in this clinically impactful process.

### Loss of *RBA1* enhances *C. auris* adhesion to plastics and human keratinocytes

A key clinical characteristic of *C. auris* is its ability to adhere to and persist on a variety of surfaces in the nosocomial environment, including medical equipment and patient skin. As our transcriptional profiling revealed loss of *RBA1* enhanced the expression of an assortment of *C. auris* adhesin-encoding genes, we hypothesized that loss of *RBA1* would enhance *C. auris* adhesion to multiple substrates. To test this, we first evaluated the adherence of our Δ*rba1* and Δ*rba1*::*RBA1^+^*strains, as well as the isogenic clinical isolates SKU013 and SKU018, to polystyrene plastic under a variety of conditions. In RPMI media, SKU018 and the Δ*rba1* strains exhibited greater adhesion than SKU013 to both polystyrene and tissue culture-treated polystyrene (**Fig. 5a, b**). This difference was magnified when the polystyrene well plates were pre-treated with human plasma, with or without the addition of 10% human serum to the media (**Fig. 5c, d**). By contrast, Δ*rba1*::*RBA1^+^*control strains phenocopied the parental SKU013 under all conditions.

**Fig 5.**
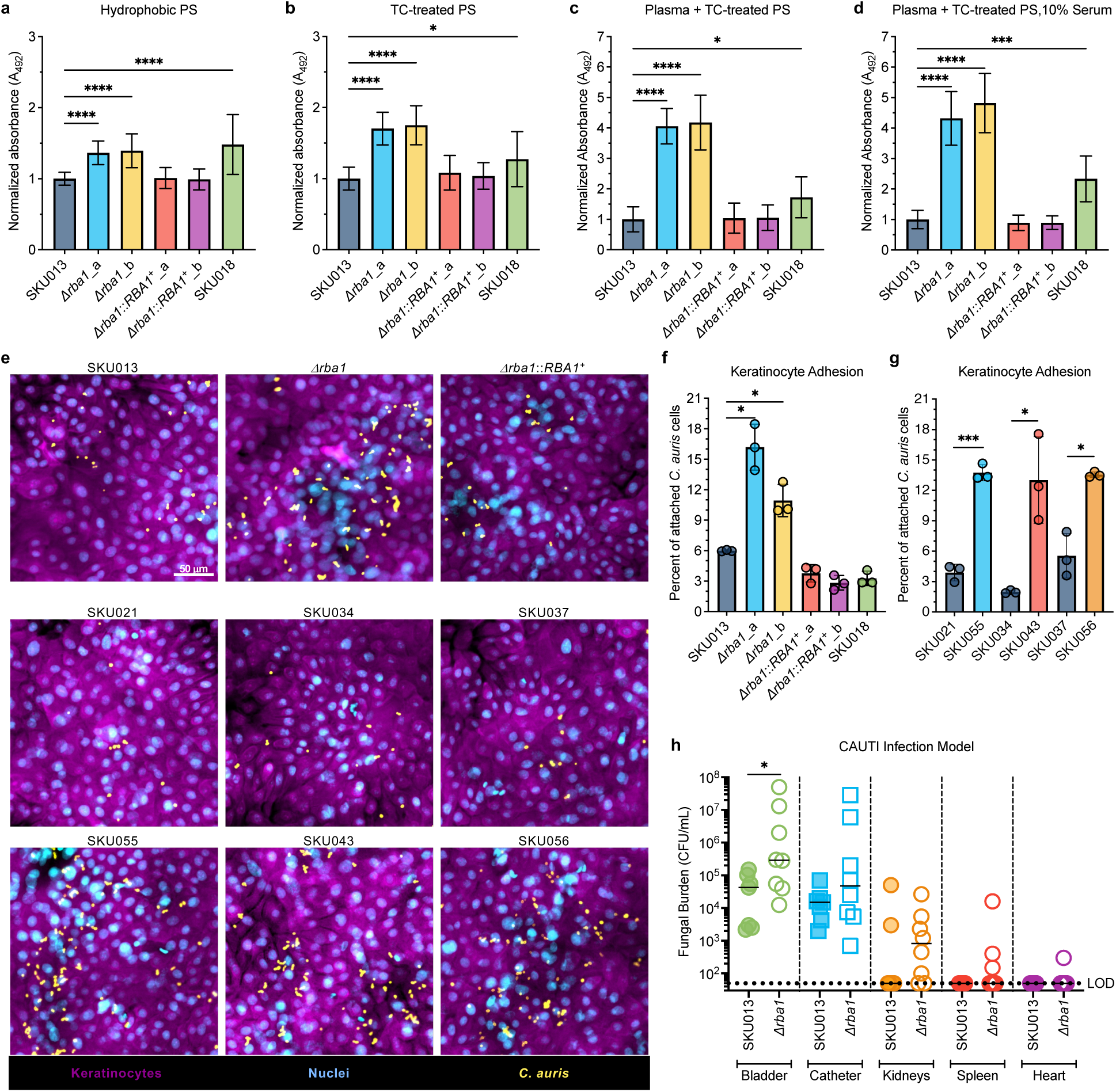
Loss of *RBA1* increases adhesion and fungal burden in murine CAUTI model. **a, b, c, d,** Adhesion of *C. auris* isolates and *RBA1* strains grown in RPMI culture media after 90 minutes to (**a**) untreated polystyrene (PS), (**b**) tissue culture (TC) -treated PS, (**c**) human plasma coated, TC-treated PS, and (**d**) human plasma coated, TC-treated PS grown in RPMI supplemented with 10% human serum. Adhesion assessed by absorbance at 490 nm following XTT treatment. Statistical significance was determined using Kruskal-Wallis one-way ANOVA tests followed by a Dunn’s test for multiple comparisons to the *RBA1*^WT^ control SKU013 (\**P* ≤ 0.05; \*\*\*\**P* < 0.0001). **e,** Representative immunofluorescence images showing adherence of *C. auris* cells to human N/TERT keratinocytes. *C. auris* was stained with FITC-conjugated anti-*Candida* antibody (yellow) and keratinocytes were stained using CellMask Deep Red (magenta) with a Hoechst counterstain for nuclei identification (blue). **f, g,** Quantification of *C. auris* isolates strains adhered to human N/TERT keratinocytes as measured by the proportion of attached cells remaining after washing. Bars represent mean ± standard deviation for three biological replicates. Statistically significant adhesion increases were determined using unpaired two-tailed Welch’s t-tests for (**e**) Δ*rba1* compared to SKU013 (\**P* ≤ 0.05) and (**g**) SKU055 (*RBA1*^Q174*^), SKU043 (*RBA1*^Q225*^), and SKU056 (*RBA1*^C303S^) compared to *RBA1*^WT^ isolates, SKU021, SKU034, and SKU037, respectively (\**P* ≤ 0.05; ***P≤0.001). **i**, Fungal burden of SKU013 and the derivative Δ*rba1* strain obtained from the catheter material or bladder, kidney, spleen, and heart tissues after infection in a murine model of catheter associated urinary tract infection (CAUTI). Two-tailed Mann-Whitney tests showed significant differences in fungal burden recovered from bladder tissue (\**P* ≤ 0.05, n=8 mice per group). CFU, colony forming units; LOD, lower limit of detection.

Next, we evaluated whether loss of *RBA1* promotes adhesion of *C. auris* to human keratinocytes. Δ*rba1* strains were observed to exhibit enhanced adherence to a confluent culture of human N/TERT keratinocytes, while neither Δ*rba1*::*RBA1^+^* controls strains nor SKU018 differed from the SKU013 control (**Fig. 5e** (top)**, f**). Outbreak isolates harboring the *RBA1*^Q174*^ (SKU055), *RBA1*^Q225*^ (SKU043), and *RBA1*^C303S^ (SKU056) mutations were similarly found to exhibit enhanced adhesion to keratinocytes as compared to their closest available genetic relatives (**Fig. 5e** (bot)**, g**) further supporting the connection between loss of *RBA1* and enhanced adherence to human keratinocytes. Taken together, these results confirm that loss of *RBA1* significantly enhances *C. auris* adhesion characteristics under a variety of conditions.

### Loss of *RBA1* increases fungal burden during catheter-associated urinary tract infections

As loss of *RBA1* increased *C. auris* adhesion, biofilm formation, and the expression of the previously characterized virulence factors *SCF1*, *IFF4109*, and *ALS4112/ALS5*, we next sought to evaluate the impact of *RBA1* on *C. auris* biofilm-related infection *in vivo.* To accomplish this, we utilized our mouse model of CAUTI ^25, 32^, where immunocompetent mice were catheterized and the bladder of each mouse was challenged with *C. auris* transurethrally 24 hours prior to sacrifice for evaluation of fungal burden and systemic dissemination of infection. Importantly, this mimics a clinical scenario where *C. auris* is often cultured from patients with urinary catheters and is relevant to multiple patient populations at increased risk of *C. auris* colonization and infection. In this model, the Δ*rba1* strain was found to exhibit a 6.8-fold increase in bladder tissue fungal burden (*p*= 0.0435) by 24 hours post-infection, as compared to SKU013 (**Fig. 5h**). Moreover, a trend toward increased dissemination of the Δ*rba1* strain to kidney and spleen tissues was also observed, and one mouse infected with the Δ*rba1* strain was found to have detectable fungal burden in cardiac tissue. These findings suggest that loss of *RBA1* may also enhance virulence of *C. auris* during clinical infections such as CAUTI.

### Mutations in *RBA1* have emerged during numerous *C. auris* clinical outbreaks

We next hypothesized that *RBA1* mutations have also likely occurred during other *C. auris* outbreaks as well. To test this, we leveraged publicly available *C. auris* WGS data collected for infection control and outbreak tracking to identify mutations in the *RBA1* gene. Ninety-four mutations in *RBA1* were identified, and 310 of 2,181 evaluable isolates were found to carry one or more of 58 identified non-synonymous or nonsense *RBA1* mutations, including *C. auris* isolates representing genetic clades I, II, and IV. The majority of the identified non-synonymous/nonsense mutations are considered as likely loss-of-function mutations (40; 61.5%) (**Fig. 6a**), similar to the mutations we initially identified from the Kuwait outbreak collection. Evaluating meta data and publications associated with identified isolates revealed 10 non-synonymous *RBA1* mutations originating from documented outbreaks of *C. auris* infection on four continents and were caused by isolates representing both genetic clades I and IV (**Fig. 6b**) ^10, 16, 33, 34, 35, 36^. Notably, this included two instances where *RBA1* mutations arose during Indian outbreaks^37^ initially reported in 2013, as well as two instances of *RBA1* mutations occurring during ongoing outbreaks in the United States^34, 38^. These findings clearly demonstrate convergent mutations in *RBA1* across multiple of *C. auris* outbreaks and further suggest that these mutations confer a competitive advantage to *C. auris* in the healthcare environment.

**Fig 6.**
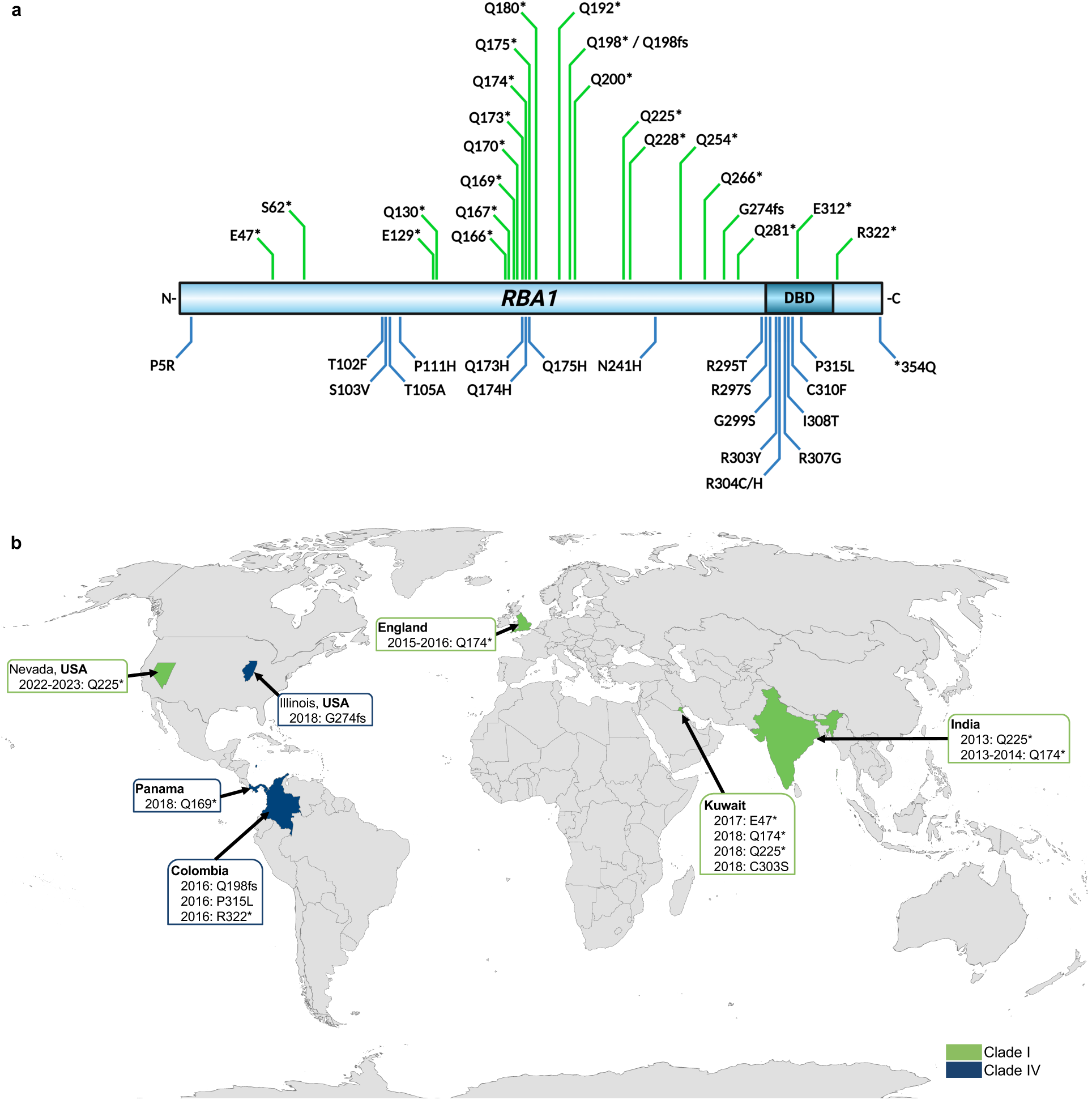
Identification of *RBA1* mutations emerging during numerous outbreaks of *C. auris* infections worldwide. **a**, Diagram depicting non-synonymous mutations in *RBA1* identified by analyzing publicly available *C. auris* whole genome sequencing data. Green lines on top of figure indicate early stop codons and frameshift mutations. Blue lines on bottom of figure indicate missense mutations. N241H and *354Q were identified among all clinical isolates classified as clade IV. DBD: DNA-binding domain. **b,** World map indicating locations of documented outbreaks of *C. auris* infections where mutations in *RBA1* were identified from publicly available data. Green fills indicate isolates representing genetic Clade I and blue fills indicate isolates representing genetic clade IV.

## DISCUSSION

*C. auris* is frequently associated with clinical outbreaks of infection and multi-drug resistance thus posing a challenge to clinicians reminiscent of the healthcare-associated bacterial pathogens, *Acinetobacter baumannii* and *Enterococcus faecium*, ^3, 8^. Unlike other agents of invasive candidiasis, *C. auris* is not considered as part of the human normal flora and is not known to be widely distributed in environmental reservoirs. Instead, *C. auris* infections are currently thought to be acquired predominantly through dissemination of recently emergent drug-resistant, and possibly infection-adapted, lineages within medical environments, leading to the infection of vulnerable patient populations^3, 16^. Thus, as with other healthcare-associated pathogens, the identification of *C. auris* evolution during clinical outbreaks and characterization of traits promoting the persistence among human patients is essential to both preventing infections and mitigating prolonged outbreaks of infection.

Here, we analyzed samples from outbreaks in Kuwait to identify genomic and microbiological elements which may promote transmission and persistence of *C. auris* in clinical settings. Rather than distinct clustering of serial isolates from individual patient cases, phylogenetic analysis identified isolate clusters originating from multiple patients indicating multiple instances of pathogen lateral transmission consistent with clinical dissemination. Interestingly, while numerous cases of isolates acquiring mutations associated with echinocandin and AmB resistance were identified, persistence and spread of these multi-drug-resistant lineages was not observed in this study. Conversely, among both phylogenetic clusters B and C, closely related isolates harboring acquired *RBA1* mutations were cultured from multiple patients over periods of time spanning many months. While there was no association between clinical antifungal susceptibility and *RBA1* genotype, *C. auris* isolates carrying clinically acquired loss-of-function mutations in *RBA1* were observed to exhibit enhanced biofilm formation as compared to those with wildtype *RBA1* sequences.

Biofilms are critically important to human public health as they promote microbial survival in a variety of environments, demonstrating increased resilience to disinfectants and antimicrobials, and contribute to pathogen dissemination and the establishment of new infections. In fact, the majority of human infections involve biofilms either within human tissues or on the surface of in-dwelling medical devices such as catheters or prosthetic implants ^39^. As numerous clinical isolates from the studied clinical outbreaks in Kuwait were isolated from the urine of catheterized patients or the respiratory secretions of patients with endotracheal tubes, we hypothesized that the identified loss-of-function mutations in *C. auris RBA1* represented a clinically emergent adaptation promoting pathogen success in the medical environment through enhanced biofilm formation. Genetic interrogation of the impact of *C. auris RBA1* definitively demonstrated loss of this gene enhanced biofilm formation when evaluated using *in vitro* models mimicking clinically relevant conditions including those specific to CAUTI. This enhancement of biofilm formation may represent a selective pressure which promoted the emergence and persistence of *RBA1* mutant lineages during the studied clinical outbreaks.

Subsequent transcriptional profiling revealed a narrow role in *C. auris* cells growing under planktonic conditions, and a much wider influence on gene expression when cells were grown in biofilms. Under both conditions, however, Δ*rba1* strains exhibited increased expression of a multitude of cell surface adhesins, including multiple genes we have previously shown to significantly impact *C. auris* virulence during infection. The two previously characterized *C. auris*-specific adhesins, *SCF1* and *ALS4112/ALS5,* were substantially elevated in the Δ*rba1* strain grown in planktonic conditions, with both genes exceeding a 7-log_2_-fold increase over the SKU013 parental control. As these adhesins are known to be critical to the ability of *C. auris* to adhere to a variety of substrates including human skin^26, 27^, this suggested that *RBA1* may impact the propensity of *C. auris* to both colonize patients and establish new biofilms. Further investigation of gene expression in biofilms demonstrated a wide variety of genes predicted to encode *C. auris* cell surface proteins, cell wall constituents, secreted enzymes, and transcriptional regulators were differentially expressed in the Δ*rba1* strain. Intriguingly, this also included the *C. auris* orthologs of *C. albicans ENG1* and *YWP1*, two secreted beta-glucanases which are known to play critical roles in cell wall beta-glucan masking^28, 29, 30^. This finding, combined with the altered expression of a multitude of GPI-anchored cell surface proteins and the differences in extracellular matrix production observed by scanning electron microscopy, suggests that the loss of *RBA1* may also facilitate evasion of *C. auris* detection from host defenses during infection.

We next examined whether loss of *RBA1* and subsequent increased expression of *SCF1* and *ALS4112/ALS5* would impact the ability of *C. auris* to adhere to surfaces where *SCF1* and *ALS4112/ALS5* play a significant role^26, 27^. Strains lacking *RBA1* were found to exhibit enhanced adhesion to human keratinocytes as well as polystyrene surfaces both with and without human plasma coatings, which mimics the deposition of host factors such as fibrinogen on medical device surfaces. Moreover, evaluation of three *C. auris* clinical isolates from the Kuwait outbreaks harboring three additional loss-of-function mutations, as compared to their respective closest available genetic relative with a *RBA1*^WT^ allele, again demonstrated loss of *RBA1* to increase adhesion of *C. auris* to human keratinocyte. This finding is of substantial clinical relevance, as it supports that loss of *RBA1* can promote the dissemination of *C. auris* and colonization of patients during a clinical outbreak. Further, patients colonized with *C. auris* strains harboring *RBA1* loss-of-function mutations may pose a greater risk of transmission within the hospital environment.

We also examined the impact of *RBA1 in vivo* using a murine CAUTI model which replicates a biofilm-related infection of a site where *C. auris* is commonly isolated. Contrary to other commonly utilized models of pathogenic yeast infection, this model uses mice that have not been immunosuppressed. Instead, immunocompetent mice are made vulnerable to *C. auris* infection through the implantation of a silicone catheter, similar to urinary catheters commonly used among human patients. In this model, the Δ*rba1* strain was found to develop a significantly greater fungal burden in the mouse bladder, and a trend towards greater dissemination of the Δ*rba1* strain from the urinary tract to other tissues including the heart was observed. These findings further suggest loss of *RBA1* is advantageous during *C. auris* clinical infection. This is clinically significant as CAUTIs often lead to urosepsis with a ∼30% 7-day mortality rate ^15, 40^. Since the urinary tract is a source of ∼25% of the all the sepsis cases, patients infected with *C. auris* strains harboring *RBA1* mutation may be predisposed to severe clinical complication.

Finally, leveraging the growing body of publicly available WGS data generated through ongoing *C. auris* infection control and prevention efforts, we were able to investigate whether mutations in *RBA1* have emerged during other clinical outbreaks. This approach uncovered dozens of distinct *RBA1* mutations, impacting hundreds of clinical isolates and more than 10% of evaluable samples. *RBA1* mutations were identified among isolates representing three of the major genetic subgroups of *C. auris,* but were notably absent among isolates from clade III, suggesting either clade-specific impacts of *RBA1* mutations or a potential bias within samples currently available in the public data set. Like the four *RBA1* mutations identified from the Kuwait outbreaks, the mutations identified from the global collection were predominantly predicted loss-of-function mutations such as those encoding early stop codons, frameshift mutations, or altering highly conserved residues in the ZCF DNA-binding domain. Identified isolates with *RBA1* mutations were found to date back to some of the earliest documented clinical outbreaks of *C. auris*, including those in India, England, and Columbia. Additionally, isolates from outbreak lineages actively disseminating in the United States were also found to harbor *RBA1* mutations. Taken together, these findings clearly demonstrate that mutations in *RBA1* have emerged repeatedly during clinical outbreaks of *C. auris* infection and strongly support the notion that the enhancement in adhesion and biofilm formation conferred by these mutations provided a competitive advantage within the medical environment in numerous previous outbreaks.

In conclusion, these results call attention to the capacity of *C. auris* to evolve and enhance traits beyond drug resistance during outbreaks of infection by highlighting the unique nature of *C. auris* among other fungal pathogens and identifying mutations in *RBA1* as a potential indicator of outbreak-adapted lineages. As traits conferred by *RBA1* mutations are advantageous to *C. auris* spread and persistence, and as we have identified numerous instances where these mutations have arisen during global outbreaks of infection, the identification and tracking of lineages carrying *RBA1* mutations may be merited as part of infection prevention efforts.

## MATERIALS AND METHODS

### Ethics Statement

*C. auris* isolates were cultured from various clinical specimens of hospitalized patients in Kuwait. The need for informed consent was waived by the Health Sciences Center Ethics Committee (Approval letter Ref: VDR/EC/3724 dated January 31, 2021) as the clinical specimens were collected as part of routine diagnostic work-up for the isolation of bacterial/fungal pathogens for suspected infections. The results are reported here on deidentified samples without revealing patient identity. All animal care was consistent with the Guide for the Care and Use of Laboratory Animals from the National Research Council. The University of Notre Dame Institutional Animal Care and Use Committee approved all mouse infections and procedures as part of protocol number 25-01-9009. For urine collection, all donors signed an informed consent form, and protocols were approved by the Institutional Review Board of the University of Notre Dame under study #19-04-5273.

### C. auris Isolates

Seventy *C. auris* isolates from previously described multi-institution outbreaks of *C. auris* infections in Kuwait, were obtained by the Mycology Reference Laboratory Department of Microbiology, Kuwait University, Kuwait ^41^. All isolates (including names used in publication of their original description) and strains used in this study are listed in **Table S1**. All *C. auris* strains were stored in 40% glycerol stocks at -80°C and cultured at 35°C in liquid YPD (1% yeast extract, 2% peptone, 2% glucose) media before plating on Sabouraud dextrose (SD) agar (BD Difco) media to ensure standardized growth for phenotypic testing.

### Plasmid and strain construction

EPIC-mediated gene editing was used to generate *RBA1* (B8441v3: B9J08_03209) modifications in *C. auris* clinical isolate SKU013 (also known as Kw1432/17). Duplexed guide oligos (Integrated DNA Technologies) (**Table S2**) targeting disruption or reversion of B9J08_03209 were added to the pJMR19 vector using restriction enzyme cloning^22^. Plasmids were maintained in *E. coli* DH5α cells cultured in Luria-Bertani broth supplemented with 5mg/L ampicillin and isolated using QIAprep® Spin Minipreps (Qiagen). Repair templates for *RBA1*-disruption and reversion were amplified from a gBlock sequence (Integrated DNA Technologies) and SKU013 gDNA, respectively, via PCR and purified with QIAquick PCR Purification Kit (Qiagen). All oligos, plasmids, and gBlocks used in this study are listed in **Table S2.** *C. auris* Lithium acetate transformations were then performed as previously described ^22^ with positive selection on 200µg/mL nourseothricin-supplemented YPD agar. *RBA1* editing was confirmed by both Sanger sequencing at the Hartwell Center at St. Jude Children’s Research Hospital (SJCRH) and with long read nanopore sequencing by Plasmidsaurus.

### Whole genome sequencing

Genomic DNA (gDNA) was extracted from 1mL of overnight liquid YPD cultures using MasterPure^TM^ Yeast DNA Purification kits (Biosearch Technologies) according to manufacturer instructions. Sample libraries were prepared from gDNA by the Hartwell Center (SJCRH) using KAPA HyperPrep (Roche) and sequencing targeting 100x coverage for paired 150bp reads was performed via Illumina NovaSeq. Paired-reads FASTQ files were analyzed using CLC genomics workbench v24.0 (QIAGEN) software. Low quality reads were trimmed with Trim Reads v2.8 and subsequently mapped to the B8441v3 genome (GCA002759435.3) according to CLC-workbench recommendations for match scores, mismatch, insertion, and deletion costs. Local Realignment v2.6 was used to refine unaligned ends and mutations were identified within annotated coding sequences via Fixed Ploidy Variant Detection v2.6 with ploidy=1.

Phylogenetic analysis was initiated with Create SNP Tree v1.9 using the Maximum Likelihood Phylogeny v1.4 algorithm. The resulting SNP phylogenetic tree was visualized from Neighbor Joining construction methods specifying Jukes Cantor nucleotide substitution modeling with estimated topology and bootstrapping for 100 replicates. Sequences were deposited in National Center for Biotechnology Information Sequence Read Archive.

### Biofilm Production Assays

To assess biofilm production in the presence of human plasma, 96-well flat-bottomed plates (VWR, 10861-562) were coated with 100 μL pooled human plasma (Innovative Research Inc, IPLAWBNAC50ML) diluted 1:5 in carbonate-bicarbonate buffer (Sigma C3041-50CAP) and incubated overnight at 4°C. Colonies from overnight SD agar plates were used to generate standardized OD_600_ = 0.5 cell suspensions^42^ in RPMI-Biofilm media, consisting of RPMI medium 1640 (Gibco) + 3-(4-Morpholino)propane sulfonic acid (MOPS, Fisher Bioreagent) + 2% dextrose, buffered to pH 7 and supplemented with 10% heat-inactivated/male AB human serum (HuS, Sigma-Aldrich). Plasma coatings were gently removed the plates and 200 μL of the cell suspensions were added to the wells and plates were sealed with a breathable (EASYSEAL) membrane. After 90-minutes at 37°C, 250 RPM, plates were gently washed sterile PBS to remove non-adhered cells. Fresh 200 μL of RPMI-Biofilm media was then added back to each well and plates were incubated for 24 hrs at 37°C, 250 RPM. For quantification of biofilm production, supernatants were removed and wells were gently washed twice with PBS. Biofilms were stained with 0.4% crystal violet as previously described^42^. Crystal violet absorbance measured at 595nm on an Agilent BioTek Synergy H1 Microplate Reader. Average media only negative controls were subtracted from the sample reads and values were normalized to SKU013.

To replicate biofilm growth replicating the urinary tract environment, 96-well flat-bottomed plates (VWR, 10861-562) were coated with 100 μl of Fg (450 μg/ml) and incubated overnight at 4°C.^25^ The various strains were grown overnight, shaking in YPD, and the inoculum was normalized to ∼1 × 10^6^ CFUs/ml. Cultures were then diluted (1:100) into human urine supplemented with 10% heat inactivated human serum and incubated in the wells of the 96-well plate at 37°C for 48 hours. Human urine from at least two healthy female donors between the ages of 20 to 35 were collected and pooled. Donors did not have a history of kidney disease, diabetes, or recent antibiotic treatment. Urine was sterilized with a 0.22-μm filter (VWR 29186-212), pH was normalized to 6.0 to 6.5, and urine was used immediately for the assays. Following biofilm formation, supernatants were removed, and plates were incubated in 200 μl of 0.5% crystal violet for 15 min. Crystal violet stain was removed, and the plate was washed with water to remove the remaining stain. Plates were dried and then incubated with 200 μl of 33% acetic acid for 15 min. In another plate, 100 μL of the acetic acid solution was transferred, and absorbance values were measured via a plate spectrophotometer at 595 nm (Molecular Devices SpectraMax ABS). Values were then normalized to SKU013.

### RNA Extraction

Normalized cell suspensions (OD_600_ = 0.1) from SD replicate platings were diluted ∼1:50 in 10 mL RPMI-Growth media, prepared from RPMI medium 1640 (Gibco) + MOPS (Fisher Bioreagent) buffered to pH 7 and grown overnight at 35°C, 220 RPM to an OD_600_ = 2.0 ± 0.3. Concentrated cultures were diluted in >20mL RPMI-Growth media to an OD_600_ = 0.2 then split into two 10mL aliquots for direct comparison of planktonic growth and biofilm development. Planktonic cells were grown over 4 hours in 50mL conical at 35°C, 220RPM and pelleted at 4000 RPM. For biofilm development cells were transferred to T75 flask pre-coated with 20% human plasma in carbonate-bicarbonate buffer prepared as described above. Following 2 hr incubation at 37°C, non-adhered cells were gently removed and fresh 10mL RPMI-Growth media was added for an additional 4 hr incubation at 37°C. Biofilms were gently washed once with PBS and biofilms were then removed by scraping in the presence of 10mL sterile PBS and pipette transfer to a 50mL conical tube for centrifuge pelleting. All pellets were flash frozen in liquid nitrogen and stored at -80°C. RNA was isolated using the Invitrogen RiboPure^TM^ Yeast kit (ThermoFisher Scientific^TM^, USA) per manufacturer’s instructions.

### RNA Sequencing

The Hartwell Center (SJCRH) prepared the RNA libraries using Illumina chemistries per manufacturer’s instructions and sequenced the RNA on an Illumina NextSeq (Illumina Inc., USA) using the total stranded RNA-Seq application for paired end runs targeting 150bp reads. Sequenced reads were analyzed with CLC Genomics Workbench version 24.0.1 (QIAGEN). Imported reads were trimmed using default settings for failed reads and adaptor sequences and then subsequently mapped to the *C. auris* B8441 reference genome (GenBank accession: GCA_002759435.3) with paired reads counted as one and expression values set to RPKM. Principal component analysis was utilized for the initial clustering assessment of biological replicates and mismatch, insertion, and deletion costs were set to default parameters for RNA-Seq Analysis 2.7. Differential expression was calculated using negative binomial GLM modeling with whole transcriptome RNA-Seq defaults for normalization with Trimmed Mean of M values, using Differential gene expression v1.4 (CLC genomics workbench, Qiagen). Comparisons were made to the wildtype *SKU013* clinical isolate for each condition. Differential expression was defined as fold changes ≥ 2 or ≤ -2 (FDR p-value ≤0.05)

### Scanning Electron Microscopy

Sterile silicon chips were loaded into 24-well plate and coated overnight with 0.5mL 20% human plasma in carbonate-bicarbonate buffer. Suspensions of SKU013, Δ*rba1*::*RBA1^+^*_a, and 11*rba1_a* were cultured in RPMI-Biofilm media as described above for biofilm production assays with 0.5mL of cells loaded into the wells. Following 24 hour incubation, wells were gently washed with 1mL PBS and biofilms were fixed with 0.1M sodium cacodylate buffer at pH 7.4 with 0.15% alcian blue, 1% tannic acid, 2.5% glutaraldehyde, 2% paraformaldehyde for 2 hours and stored at 4°C. Preparation, staining, and imaging the fixed biofilms was performed by the Electron Microscopy Division of the Cell and Tissue Imaging Center (CTIC-EM) at SJCRH for SEM Fixed biofilms were washed with 0.1M sodium cacodylate buffer + 0.15% alcian blue prior to osmium tetroxide staining in the presence of 0.15% alcian blue. Imaging was performed on a Zeiss GeminiSEM 460. Scale bars were embedded using ImageJ^43^.

### Polystyrene adhesion assay

Adhesion to flat-bottom 96-well non-tissue culture treated polystyrene plates (CS100-701311, NEST SCIENTIFIC) was assessed in RPMI-Growth+ media, defined as RPMI medium 1640 + MOPS + 2% dextrose, buffered to pH 7. Adhesion to tissue culture treated plates (TP92696, TPP Midwest Scientific), was assessed for RPMI-Growth+ media and RPMI-Biofilm media in the presence and absence of human plasma, coated as described above. cells cultured and plates were loaded as described for biofilm production assays. Following a 90-minute incubation at 37°C, 250RPM, wells were gently washed once with PBS to remove non-adhered cells. Adhered cells were stained with 100μL of a 10:1 solution of XTT sodium salt and phenazine methosulfate (VWR International LLC). Absorbances at OD_492_ and OD_630_ were read after 60 minutes and values were normalized to SKU013.

### Keratinocyte adhesion assay

Adhesion of *C. auris* to human keratinocytes was determined as previously described^26^. Human N/TERT keratinocytes^44^ were grown in 96-well plates to complete confluence in keratinocyte-SFM medium (Gibco, 10724-011) supplemented with 30 µg/ml bovine pituitary extract, 0.2 ng/ml epidermal growth factor, and 0.31 mM CaCl_2_. Overnight *C. auris* cultures were diluted to an OD_600_ of 0.03, and 100 µl of this suspension was added to each well containing keratinocytes. In parallel, an equal volume of culture was added to empty wells, centrifuged (210 g, 1 min), and imaged to determine input cell counts, while plates containing keratinocytes were centrifuged under the same conditions and incubated at 37 °C with 5% CO_2_ for 1 h. Non-adherent yeast cells were then removed by washing three times with PBS. The remaining cells were fixed in 4% formaldehyde and stained with FITC-conjugated anti-*Candida* antibody (Meridian Bioscience, B65411F; 1:1000). Keratinocyte nuclei and cell bodies were labeled with Hoechst (Thermo Fisher, 62249; 1:1000) and CellMask Deep Red (Thermo Fisher, H32721; 1:5000), respectively.

Fluorescent images were acquired at 10X magnification using a BioTek Lionheart FX microscope. The number of adherent *C. auris* cells was quantified using a customized CellProfiler analysis pipeline^26, 45^. Numbers of input cells were determined from brightfield images pre-processed in Fiji/ImageJ (v2.14.0)^43^ using a custom macro for edge-based segmentation and binary mask generation. The proportion of adherent to input cells was calculated to assess the adhesion efficiency of each *C. auris* strain.

### Catheter-Associated Urinary Tract Infection (CAUTI) Modeling

Female C57BL/6 mice (Jackson Laboratory) were subjected to transurethral implantation of a silicone catheter followed by transurethral inoculation of the bladder lumen with ∼5 × 10^5^ colony forming units (CFU) in PBS, as previously described ^32^. Mice were euthanized at 24 hours post-infection by cervical dislocation after anesthesia inhalation and the catheter, bladder, kidneys, spleen, and heart were aseptically harvested. Bladders were weighed and then homogenized for CFUs. The other organs were homogenized, and catheters were cut into small pieces before sonication for fungal CFU enumeration.

### Statistics and Reproducibility

Statistical parameters and applied analysis methods including exact sample numbers (n) and definitions for representative central value and spread (mean ± SEM, CFU quantifications; representative modal value (n=3), minimum inhibitory concentrations) are annotated in corresponding figure legends. Data from at least three experiments were pooled for each adhesion and biofilm assay and twice for each *in vivo* mouse assay. One-way ANOVA tests, two-tailed Welch’s t-tests, and two-tailed Mann-Whitney *U* tests were performed with GraphPad Prism 10 software (GraphPad Software) for comparisons of described in-vitro assays and in vivo CAUTI experiments, respectively.

For RNA sequencing, statistical analysis was performed using CLC Genomics Workbench version 20.0 (QIAGEN) with paired reads counted as one and expression values set to RPKM. Whole transcriptome differential gene expression analysis was performed with mismatch, insertion, and deletion costs set to default parameters, and a Wald test was used to compare SKU013 against Δ*rba1_a*. Differential expression determined by a fold change of ≥ 2 or ≤ -2 with a False discovery rate (FDR) corrected p value ≤0.05.

## ACKNOWLEDGMENTS

This work was supported in part by the St. Jude Children’s Research Hospital Children’s Infection Defense Center and Center for Infectious Diseases Research awards granted to JMR, by the National Institutes of Health (NIH)/ National Institute of Allergy and Infectious Diseases (NIAID) grant U19-AI181767 awarded to TRP, and by the NIH/ NIAID F32 AI181164 awarded to GZ. This work was supported by institutional funds from the University of Notre Dame (to FHST and ALFM), and by grants from the Good Venture Foundation (Open Philanthropy) (to ALFM), from the National Institutes of Health R01AI177875 and R21AI171742 (to FHST), and R01DK128805 (to ALFM). This research included experiments conducted by the Hartwell Center for Bioinformatics & Biotechnology, the St. Jude Center for Advanced Genome Engineering and the Cell and Tissue Imaging Center - Electron Microscopy (CTIC-EM) which are supported in part by ALSAC and the National Cancer Institute at the National Institutes of Health [P30 CA021765]. The funders had no role in study design, data collection and interpretation, or the decision to submit the work for publication.

## AUTHOR CONTIBUTIONS

Conceptualization, J.M.R., T.R.O., F.H.S.T., and A.L.F.M.; methodology, J.M.R., L.A.D., A.A.L., S.J.J., Q.Z., T.R.O., and A.L.F.M.; Investigation, L.A.D., A.A.L., S.J.J., Q.Z., Q.J.C., and J.M.R.; writing—original draft, L.A.D. and J.M.R.; writing—review & editing, L.A.D., A.A.L., S.J.J., G.Z., M.A., Q.J.C., S.A., E.M., I.A., W.A., K.A., T.R.O., F.H.S.T., A.L.F.M., and J.M.R.; funding acquisition J.M.R., T.R.O., F.H.S.T., A.L.F.M., and Q.Z.; supervision, T.R.O., F.H.S.T., A.L.F.M., and J.M.R..

## SUPPLEMENTAL DATA

### Patient 51 and 53 Case Descriptions

Patient 51 was a 54-year-old male with a past medical history significant for multiple myeloma admitted to the intensive care unit (ICU) of Hospital F, a tertiary care center associated with Kuwait Cancer Control Center in Kuwait which cares for immunocompromised patients including cancer patients, burn patients and transplant recipients diagnosed and requiring treatment for infectious diseases. Patient 51 was receiving a 14-day course of empiric caspofungin treatment when *C. auris* isolate SKU013 (*RBA1*^WT^; wildtype-*RBA1*) was first identified from a culture of their endotracheal tube and considered to represent colonization (**Fig. 2a**). Four successive isolates (**Fig. 2c**) were identified from endotracheal tube cultures, and the presence of respiratory symptoms prompted a 21-day course of empiric caspofungin treatment. During this treatment, isolate SKU018 (*RBA1*^E47*^) was obtained from the patient’s urinary catheter. SKU018 notably exhibited *in vitro* resistance to echinocandins (*FKS1*^S639F^), although it was still thought to represent likely colonization. SKU020 was obtained from a blood culture the day before patient 51 expired. SKU020 also exhibited elevated *in vitro* echinocandin resistance and differed from the earlier respiratory tract isolates SKU016 and SKU017 by only one coding region SNP (*FKS1*^R1354H^), indicating likely dissemination from the respiratory tract.

Patient 53 was a 33-year-old female in the ICU of the same tertiary care hospital (Hospital F) with a significant past medical history for Hodgkin’s lymphoma and advanced stage nodular sclerosis. Approximately one month after receiving a 21-day empiric course of liposomal amphotericin B (AmB) for suspected respiratory infection, the patient’s respiratory condition worsened and *C. auris* (SKU022) was isolated from the patient’s endotracheal tube. Notably, isolate SKU022 harbored the same *RBA1*^E47*^ mutation present in the isolate previously cultured from the urine of patient 51 (SKU018), and genetically these two isolates differed by four coding region SNPs, suggestive of pathogen lateral transmission. Following completion of a 14-day course of caspofungin treatment, two subsequent *C. auris* isolates (both *RBA1*^E47*^) were cultured from endotracheal tube and tracheal aspirate cultures, prompting a second 14-day course of liposomal AmB. After this second course of AmB, all cultures from respiratory sources remained negative. However, one month later the patient was suspected to have developed a CAUTI and urine cultures yielded isolate SKU031 (*RBA1*^E47*^). A second 14-day course of caspofungin was initiated after SKU031 was found to be AmB-resistant. Subsequently, AmB-susceptible *C. auris* (SKU032) was cultured from the patient’s urine and a third course of liposomal AmB was given. During this third course of AmB, two additional isolates were obtained from urine cultures, SKU033 (*RBA1*^E47*^) and SKU035 (*RBA1*^E47*^), and exhibited decreased susceptibility to AmB and echinocandins, respectively. Patient 53 was then transferred to another care facility, precluding further follow-up.

**Supplemental Table 1.** *Candidozyma auris* isolates and strains

| CLINICAL ISOLATES |  |  |  |  |  |
| --- | --- | --- | --- | --- | --- |
| SKU_ID | Mycology<br>Reference<br>Laboratory ID | Patient<br>ID | Subclade | Isolation source | Previous<br>citations |
| SKU001 | Kw1983-16 | 47 | lc | Tissue | 46 |
| SKU002 | Kw2031-16 | 47 | lc | Ab. drain fluid | 46 |
| SKU003 | Kw2236-16 | 47 | lc | Pus | 46 |
| SKU004 | Kw2254-16 | 47 | lc | Tissue | 46, 24 |
| SKU005 | Kw2442-16 | 30 | lc | Urine | 23, 46 |
| SKU006 | Kw2515-16 | 30 | lc | Tracheal sec. | 23, 46 |
| SKU007 | Kw2555-16 | 30 | lc | Urine | 23, 46 |
| SKU008 | Kw369-17 | 30 | lc | Tracheal sec. | 23, 46 |
| SKU009 | Kw1138-17 | 30 | lc | Urine | 23, 46 |
| SKU010 | Kw1239-17 | 30 | lc | Urine | 23, 46 |
| SKU011 | Kw1308-17 | 30 | lc | Urine | 23, 46 |
| SKU012 | Kw1388-17 | 30 | lc | Tracheal sec. | 23, 46 |
| SKU013 | Kw1432-17 | 51 | lb | ETT | 46, 24, 23 |
| SKU014 | Kw1695-17 | 51 | lb | Tracheal aspirate | 46, 24 |
| SKU015 | Kw1825-17 | 51 | lb | Tracheal aspirate | 46, 24 |
| SKU016 | Kw1856-17 | 51 | lb | Tracheal aspirate | 46, 24 |
| SKU017 | Kw1893-17 | 51 | lb | ETT | 46, 24, 23 |
| SKU018 | Kw1994-17 | 51 | lb | Urine | 46, 24, 23 |
| SKU019 | Kw2040-17 | 49 | lb | Vaginal swab | 46, 24 |
| SKU020 | Kw2074-17 | 51 | lb | Blood | 46, 24, 23 |
| SKU021 | Kw2260-17 | 52 | lb | Blood | 46, 24, 23 |
| SKU022 | Kw2501-17 | 53 | lb | ET aspirate | 46, 24, 23 |
| SKU023 | Kw2647-17 | 53 | lb | ET aspirate | 46, 24 |
| SKU024 | Kw2713-17 | 52 | lb | Urine | 46, 24, 23 |
| SKU025 | Kw2742-17 | 52 | lb | Urine | 24, 23 |
| SKU026 | Kw2871-17 | 53 | lb | Tracheal sec. | 24 |
| SKU027 | Kw2968-17 | 57 | lb | Urine | 24 |
| SKU028 | Kw2989-17 | 57 | lb | ETT | 24 |
| SKU029 | Kw3092-17 | 57 | lb | Catheter tip | 24, 23 |
| SKU030 | Kw3134-17 | 57 | lb | Urine | 24, 23 |
| SKU031 | Kw3135-17 | 53 | lb | Urine | 24, 23 |
| SKU032 | Kw3367-17 | 53 | lb | Urine | 24 |
| SKU033 | Kw3424-17 | 53 | lb | Urine | 24, 23 |
| SKU034 | Kw3506-17 | 59 | lc | Urine | 47 |
| SKU035 | Kw3521-17 | 53 | lb | Urine | 24, 23 |
| SKU036 | Kw3525-17 | 59 | lc | Tracheal sec. | 47 |

| SKU037 | Kw3584-17 | 59 | lc | Urine | 47 |
| --- | --- | --- | --- | --- | --- |
| SKU038 | Kw55-18 | 65 | lb | Sputum | 24, 23 |
| SKU039 | Kw60-18 | 59 | lc | Tracheal sec. | 23, 47 |
| SKU040 | Kw87-18 | 59 | lc | Blood | 23, 47 |
| SKU041 | Kw93-18 | 59 | lc | Tracheal sec. | 23, 47 |
| SKU042 | Kw108-18 | 59 | lc | Tracheal sec. | 23, 47 |
| SKU043 | Kw137-18 | 64 | lc | Blood | 41 |
| SKU044 | Kw147-18 | 65 | lb | Blood | 24, 23 |
| SKU045 | Kw257-18 | 68 | lc | Urine | 41 |
| SKU046 | Kw334-18 | 65 | lb | Oral swab | 24, 23 |
| SKU047 | Kw532-18 | 71 | lc | Urine | 41 |
| SKU048 | Kw1014-18 | 79 | lc | Blood | 41 |
| SKU049 | Kw1148-18 | 80 | lc | Urine | 41 |
| SKU050 | Kw1201-18 | 77 | lc | Bronchial washing | 41 |
| SKU051 | Kw1252-18 | 81 | lc | Blood | 41 |
| SKU052 | Kw1517-18 | 68 | lc | Urine | 41 |
| SKU053 | Kw1610-18 | 84 | lc | Urine | 41 |
| SKU054 | Kw1674-18 | 89 | lb | Urine | 24 |
| SKU055 | Kw1753-18 | 89 | lb | Blood | 24 |
| SKU056 | Kw2103-18 | 92 | lc | Urine | 41 |
| SKU057 | Kw2201-18 | 100 | lb | Urine | 41 |
| SKU057 | Kw2201-18 | 100 | lb | Urine | 41 |
| SKU058 | Kw2272-18 | 102 | lb | Blood | 41 |
| SKU059 | Kw2427-18 | 103 | lb | Blood | 48 |
| SKU060 | Kw2466-18 | 105 | lc | Blood | 41 |
| SKU061 | Kw2518-18 | 101 | lb | Urine | 41 |
| SKU062 | Kw2570-18 | 100 | lb | Urine | 41 |
| SKU063 | Kw2717-18 | 104 | lc | Tracheal sec. | 41 |
| SKU064 | Kw2821-18 | 100 | lb | Urine | 41 |
| SKU065 | Kw2885-18 | 100 | lb | Blood | 41 |
| SKU066 | Kw2898-18 | 113 | lc | Urine | 41 |
| SKU067 | Kw2999-18 | 113 | lc | Urine | 41 |
| SKU068 | Kw3014-18 | 118 | lb | Urine | 41 |
| SKU069 | Kw3144-18 | 125 | lc | Blood | 41 |
| SKU070 | Kw3204-18 | 123 | lc | Groin swab | 41 |
| Strain ID | Background | *RBA1* Genotype |  | Citations |  |
| *Δrba1_a* | SKU013 | B9J08_003209:c.950CAGT>CTTAT |  | This study |  |
| *Δrba1_b* | SKU013 | B9J08_003209:c.950CAGT>CTTAT |  | This study |  |
| *Δrba1::RBA1<sup>+</sup>_a* | *Δrba1_b* | *RBA1*<sup>WT</sup> |  | This study |  |
| *Δrba1::RBA1<sup>+</sup>_b* | *Δrba1_a* | *RBA1*<sup>WT</sup> |  | This study |  |

**Supplemental Table 2.**
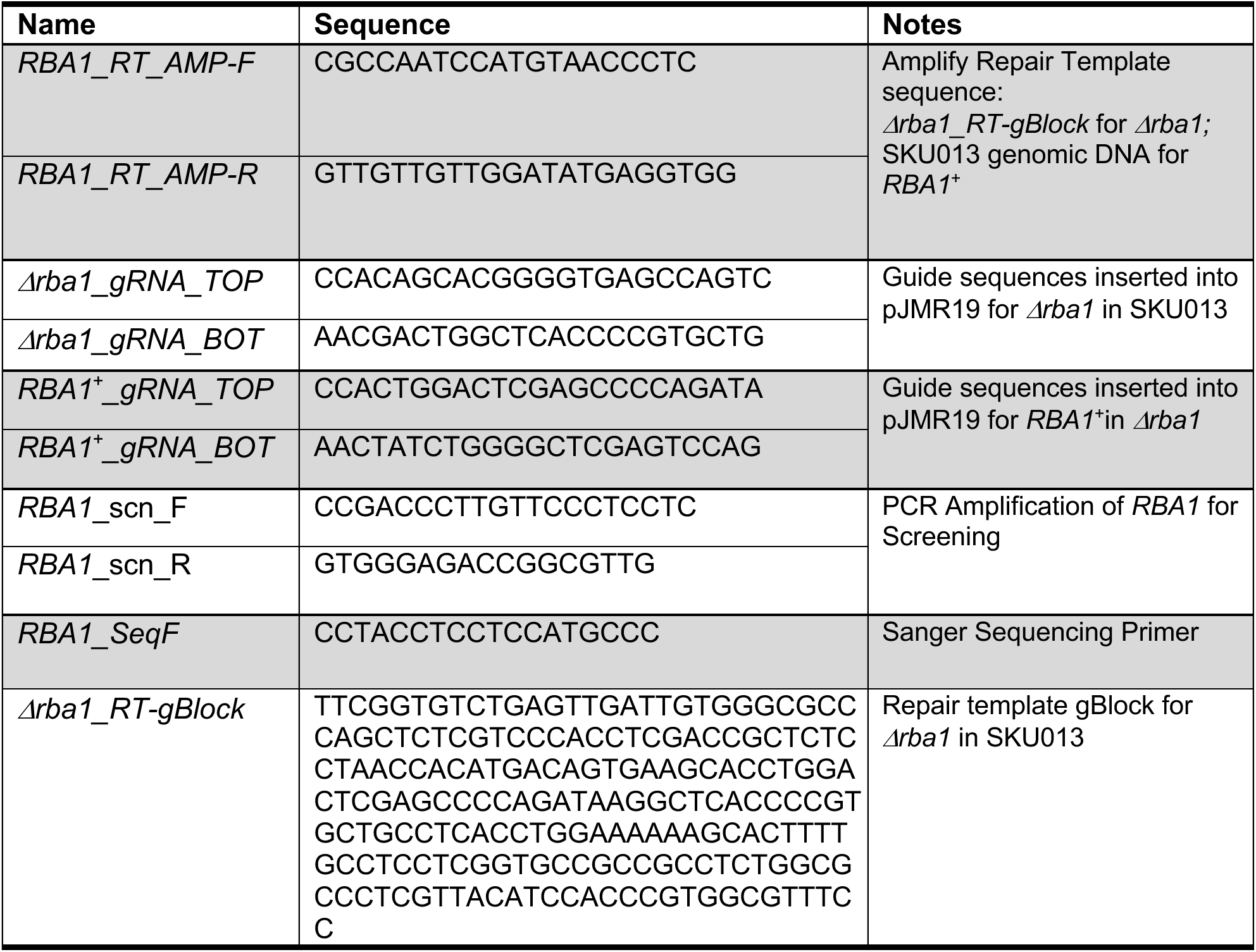
Oligos and Primers

**Supplementary Figure 1.**
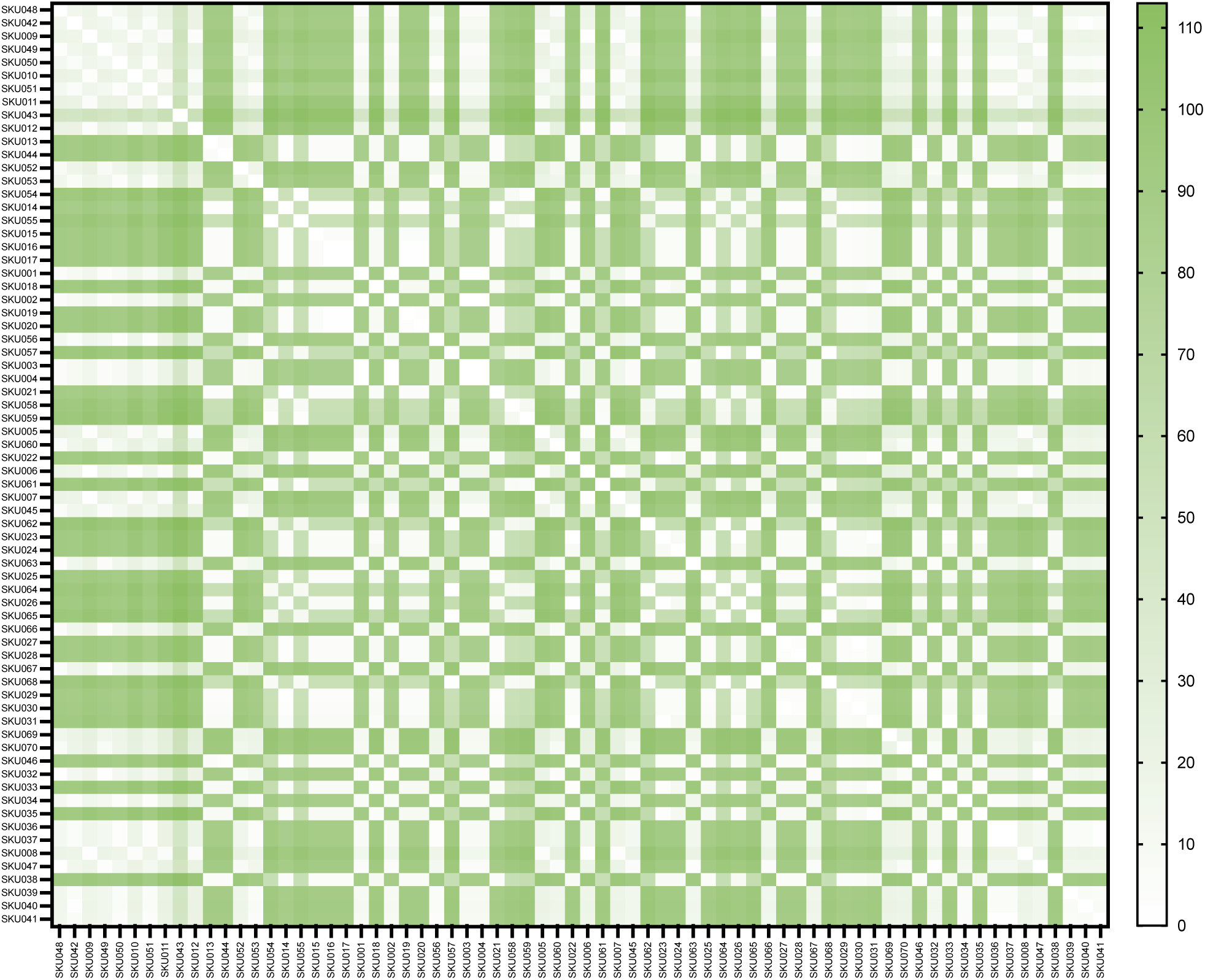
Matrix of identified single nucleotide polymorphisms (SNP) in coding regions of genome. *C. auris* isolate Genomes aligned to reference genome GCA002759435.3 (clade I, B8441_v3) for SNP identifications. SNP calling performed with fixed ploidy variant detection in CLC genomics workbench (Qiagen). 46,231 SNPs identified among the annotated open reading frame with identification of 46,055 SNPs leading to amino acid changes across all isolates.

