## Supplemental Information for "Convergent evolution on *RBA1* regulates *Candidozyma auris* host adaptation"

**Emergence and spread of outbreak-adapted *Candidozyma auris* harboring mutations in *RBA1***

**Supplemental Table 1. *Candidozyma auris* isolates and strains**

**CLINICAL ISOLATES**

| SKU_ID | Mycology Reference Laboratory ID | Patient ID | Subclade | Isolation source | Previous citations |
|---|---|---|---|---|---|
| SKU001 | Kw1983-16 | 47 | Ic | Tissue | [1] |
| SKU002 | Kw2031-16 | 47 | Ic | Ab. drain fluid | [1] |
| SKU003 | Kw2236-16 | 47 | Ic | Pus | [1] |
| SKU004 | Kw2254-16 | 47 | Ic | Tissue | [1], [2] |
| SKU005 | Kw2442-16 | 30 | Ic | Urine | [3], [1] |
| SKU006 | Kw2515-16 | 30 | Ic | Tracheal sec. | [3], [1] |
| SKU007 | Kw2555-16 | 30 | Ic | Urine | [3], [1] |
| SKU008 | Kw369-17 | 30 | Ic | Tracheal sec. | [3], [1] |
| SKU009 | Kw1138-17 | 30 | Ic | Urine | [3], [1] |
| SKU010 | Kw1239-17 | 30 | Ic | Urine | [3], [1] |
| SKU011 | Kw1308-17 | 30 | Ic | Urine | [3], [1] |
| SKU012 | Kw1388-17 | 30 | Ic | Tracheal sec. | [3], [1] |
| SKU013 | Kw1432-17 | 51 | Ib | ETT | [1], [2], [3] |
| SKU014 | Kw1695-17 | 51 | Ib | Tracheal aspirate | [1], [2] |
| SKU015 | Kw1825-17 | 51 | Ib | Tracheal aspirate | [1], [2] |
| SKU016 | Kw1856-17 | 51 | Ib | Tracheal aspirate | [1], [2] |
| SKU017 | Kw1893-17 | 51 | Ib | ETT | [1], [2], [3] |
| SKU018 | Kw1994-17 | 51 | Ib | Urine | [1], [2], [3] |
| SKU019 | Kw2040-17 | 49 | Ib | Vaginal swab | [1], [2] |
| SKU020 | Kw2074-17 | 51 | Ib | Blood | [1], [2], [3] |
| SKU021 | Kw2260-17 | 52 | Ib | Blood | [1], [2], [3] |
| SKU022 | Kw2501-17 | 53 | Ib | ET aspirate | [1], [2], [3] |
| SKU023 | Kw2647-17 | 53 | Ib | ET aspirate | [1], [2] |
| SKU024 | Kw2713-17 | 52 | Ib | Urine | [1], [2], [3] |
| SKU025 | Kw2742-17 | 52 | Ib | Urine | [2], [3] |
| SKU026 | Kw2871-17 | 53 | Ib | Tracheal sec. | [2] |
| SKU027 | Kw2968-17 | 57 | Ib | Urine | [2] |
| SKU028 | Kw2989-17 | 57 | Ib | ETT | [2] |
| SKU029 | Kw3092-17 | 57 | Ib | Catheter tip | [2], [3] |
| SKU030 | Kw3134-17 | 57 | Ib | Urine | [2], [3] |
| SKU031 | Kw3135-17 | 53 | Ib | Urine | [2], [3] |
| SKU032 | Kw3367-17 | 53 | Ib | Urine | [2] |
| SKU033 | Kw3424-17 | 53 | Ib | Urine | [2], [3] |
| SKU034 | Kw3506-17 | 59 | Ic | Urine | [4] |
| SKU035 | Kw3521-17 | 53 | Ib | Urine | [2], [3] |
| SKU036 | Kw3525-17 | 59 | Ic | Tracheal sec. | [4] |

| | | | | | |
|---|---|---|---|---|---|
| SKU037 | Kw3584-17 | 59 | Ic | Urine | 4 |
| SKU038 | Kw55-18 | 65 | Ib | Sputum | 2, 3 |
| SKU039 | Kw60-18 | 59 | Ic | Tracheal sec. | 3, 4 |
| SKU040 | Kw87-18 | 59 | Ic | Blood | 3, 4 |
| SKU041 | Kw93-18 | 59 | Ic | Tracheal sec. | 3, 4 |
| SKU042 | Kw108-18 | 59 | Ic | Tracheal sec. | 3, 4 |
| SKU043 | Kw137-18 | 64 | Ic | Blood | 5 |
| SKU044 | Kw147-18 | 65 | Ib | Blood | 2, 3 |
| SKU045 | Kw257-18 | 68 | Ic | Urine | 5 |
| SKU046 | Kw334-18 | 65 | Ib | Oral swab | 2, 3 |
| SKU047 | Kw532-18 | 71 | Ic | Urine | 5 |
| SKU048 | Kw1014-18 | 79 | Ic | Blood | 5 |
| SKU049 | Kw1148-18 | 80 | Ic | Urine | 5 |
| SKU050 | Kw1201-18 | 77 | Ic | Bronchial washing | 5 |
| SKU051 | Kw1252-18 | 81 | Ic | Blood | 5 |
| SKU052 | Kw1517-18 | 68 | Ic | Urine | 5 |
| SKU053 | Kw1610-18 | 84 | Ic | Urine | 5 |
| SKU054 | Kw1674-18 | 89 | Ib | Urine | 2 |
| SKU055 | Kw1753-18 | 89 | Ib | Blood | 2 |
| SKU056 | Kw2103-18 | 92 | Ic | Urine | 5 |
| SKU057 | Kw2201-18 | 100 | Ib | Urine | 5 |
| SKU057 | Kw2201-18 | 100 | Ib | Urine | 5 |
| SKU058 | Kw2272-18 | 102 | Ib | Blood | 5 |
| SKU059 | Kw2427-18 | 103 | Ib | Blood | 6 |
| SKU060 | Kw2466-18 | 105 | Ic | Blood | 5 |
| SKU061 | Kw2518-18 | 101 | Ib | Urine | 5 |
| SKU062 | Kw2570-18 | 100 | Ib | Urine | 5 |
| SKU063 | Kw2717-18 | 104 | Ic | Tracheal sec. | 5 |
| SKU064 | Kw2821-18 | 100 | Ib | Urine | 5 |
| SKU065 | Kw2885-18 | 100 | Ib | Blood | 5 |
| SKU066 | Kw2898-18 | 113 | Ic | Urine | 5 |
| SKU067 | Kw2999-18 | 113 | Ic | Urine | 5 |
| SKU068 | Kw3014-18 | 118 | Ib | Urine | 5 |
| SKU069 | Kw3144-18 | 125 | Ic | Blood | 5 |
| SKU070 | Kw3204-18 | 123 | Ic | Groin swab | 5 |

**STRAINS**

| Strain ID | Background | *RBA1* Genotype | Citations |
|---|---|---|---|
| *Δrba1*_a | SKU013 | B9J08_003209:c.950CAGT>CTTAT | This study |
| *Δrba1*_b | SKU013 | B9J08_003209:c.950CAGT>CTTAT | This study |
| $\Delta rba1::RBA1^{+}$_a | *Δrba1*_b | $RBA1^{WT}$ | This study |
| $\Delta rba1::RBA1^{+}$_b | *Δrba1*_a | $RBA1^{WT}$ | This study |

St. Jude - Confidential

**Supplemental Table 2. Oligos and Primers**

| Name | Sequence | Notes |
|---|---|---|
| *RBA1_RT_AMP-F*<br>*RBA1_RT_AMP-R* | CGCCAATCCATGTAACCCTC<br>GTTGTTGTTGGATATGAGGTGG | Amplify Repair Template sequence: *Δrba1_RT-gBlock* for *Δrba1;* SKU013 genomic DNA for *RBA1*$^+$ |
| *Δrba1_gRNA_TOP*<br>*Δrba1_gRNA_BOT* | CCACAGCACGGGGTGAGCCAGTC<br>AACGACTGGCTCACCCCGTGCTG | Guide sequences inserted into pJMR19 for *Δrba1* in SKU013 |
| *RBA1*$^+$*_gRNA_TOP*<br>*RBA1*$^+$*_gRNA_BOT* | CCACTGGACTCGAGCCCCAGATA<br>AACTATCTGGGGCTCGAGTCCAG | Guide sequences inserted into pJMR19 for *RBA1*$^+$in *Δrba1* |
| *RBA1*_scn_F<br>*RBA1*_scn_R | CCGACCCTTGTTCCCTCCTC<br>GTGGGAGACCGGCGTTG | PCR Amplification of *RBA1* for Screening |
| *RBA1_SeqF* | CCTACCTCCTCCATGCCC | Sanger Sequencing Primer |
| *Δrba1_RT-gBlock* | TTCGGTGTCTGAGTTGATTGTGGGCGCCCAGCTCTCG TCCCACCTCGACCGCTCTCCTAACCACATGACAGTGA AGCACCTGGACTCGAGCCCCAGATAAGGCTCACCCCG TGCTGCCTCACCTGGAAAAAAGCACTTTTGCCTCCTC GGTGCCGCCGCCTCTGGCGCCCTCGTTACATCCACCC GTGGCGTTTCC | Repair template gBlock for *Δrba1* in SKU013 |

**Supplementary Figure 1. Matrix of single nucleotide polymorphisms in coding regions identified during phylogenetic analysis**

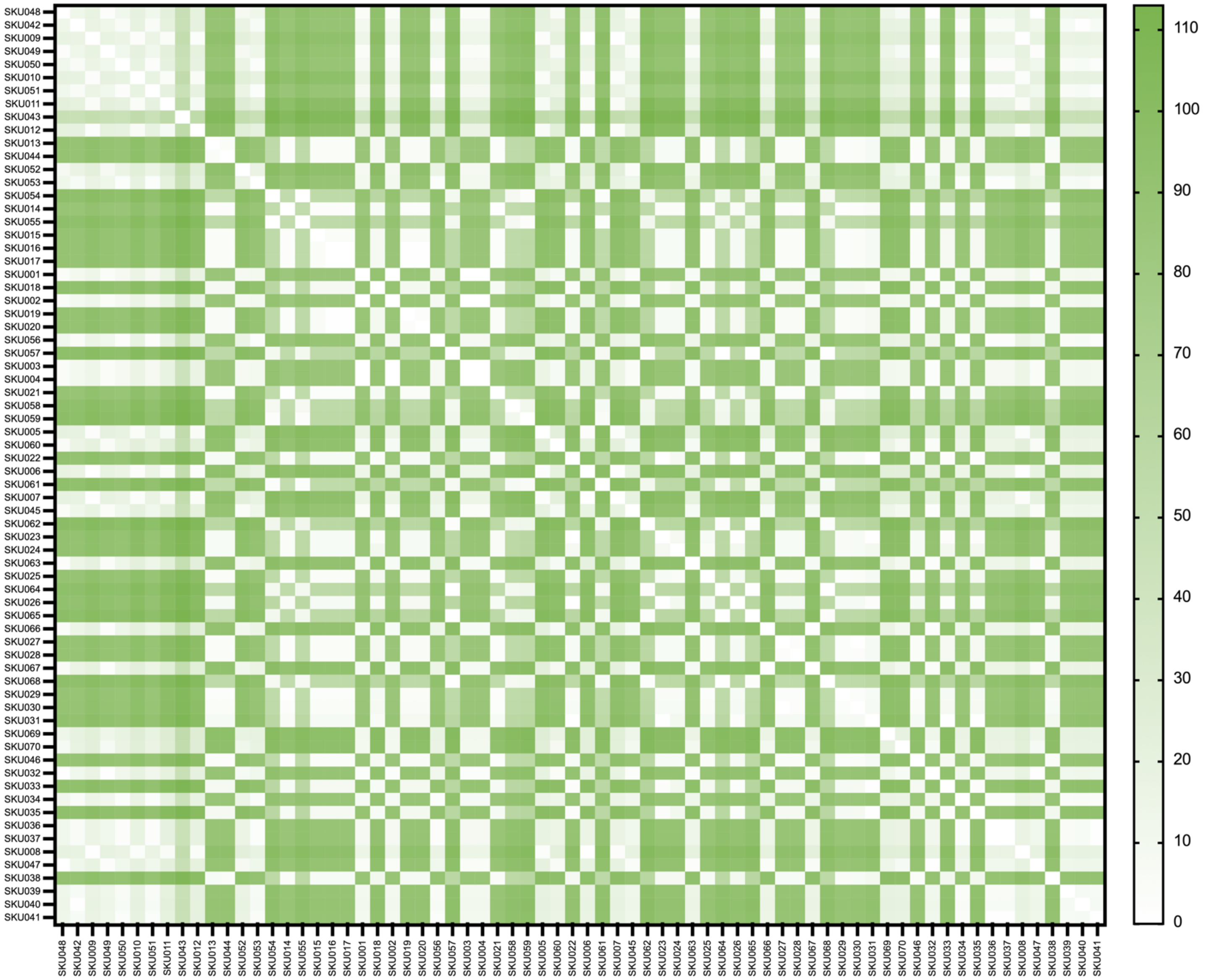
